# Interpretable spatial cell learning enhances the characterization of patient tissue microenvironments with highly multiplexed imaging data

**DOI:** 10.1101/2023.03.26.534306

**Authors:** Peng Lu, Karolyn A. Oetjen, Stephen T. Oh, Daniel L.J. Thorek

## Abstract

Multiplexed imaging technologies enable highly resolved spatial characterization of cellular environments. However, exploiting these rich spatial cell datasets for biological insight is a considerable analytical challenge. In particular, effective approaches to define disease-specific microenvironments on the basis of clinical outcomes is a complex problem with immediate pathological value. Here we present InterSTELLAR, a geometric deep learning framework for multiplexed imaging data, to directly link tissue subtypes with corresponding cell communities that have clinical relevance. Using a publicly available breast cancer imaging mass cytometry dataset, InterSTELLAR allows simultaneous tissue type prediction and interested community detection, with improved performance over conventional methods. Downstream analyses demonstrate InterSTELLAR is able to capture specific pathological features from different clinical cancer subtypes. The method is able to reveal potential relationships between these regions and patient prognosis. InterSTELLAR represents an application of geometric deep learning with direct benefits for extracting enhanced microenvironment characterization for multiplexed imaging of patient samples.

## 1 Introduction

Disease states are associated with a complex interplay of diverse cell types within close proximity in heterogeneous tissues. Highly multiplexed imaging techniques now enable the concurrent quantification of at least 40 antigens from histological specimens at subcellular resolution *in situ* [1,2]. This provides a means to assess tissue microenvironments with both molecularly specific features and cell location information. Currently, sophisticated multiplexed imaging protocols are being developed, including imaging mass cytometry (IMC) [3–6], co-detection by indexing (CODEX) [7, 8], cyclic immunofluorescence (CyCIF) [9] and multiplexed ion beam imaging (MIBI) [10, 11].

With rich cellular and neighbourhood information captured, proper analysis of these spatial data has become a new challenge. Traditional analysis methods cluster cells into distinct communities using unsupervised machine learning algorithms, on the basis of the cell-type composition mixtures of their neighbours [3, 4, 8]. However, these strategies overlook spatial inter-cellular relationships from tissues with different topological structures; hence, they can only provide a highly resolved view of cellular heterogeneity in tissues. This limits higher-order cellular community identification, such as detection of disease relevant areas.

To overcome this challenge, there has been increased interest in applying graph neural networks (GNN) [12, 13] to spatial cell analysis, in which both cell marker expressions and spatial information are taken into consideration [14–20]. Some methods focus only on tissue-scale classification [14] or cell phenotype annotation [19]; other methods have been developed for microenvironment analysis [15, 16]; and finally, some are directed at general tissue structure classification, through integrating GNN and unsupervised learning algorithms [17, 18]. However, these methods solely focus on either patient-level outcome [14] or cell-scale analysis [15–19].

Connecting the emerging single-cell rich information with spatially relevant contextual information is a challenge – specifically as it relates to correlating outcomes and cell communities of interest. SPACE-GM [20] solved this issue by training a GNN with tissue-scale labels, then combining trained latent features and K-means clustering to identify disease relevant microenvironments. Though powerful, this framework requires downstream unsupervised clustering to locate potential interested communities related to the corresponding tissue-level outcomes.

In this work, we present **Inter**pretable **S**pa**T**ial c**ELL** lea**AR**ning (InterSTELLAR), a geometric deep learning framework for multiplexed imaging data, to link the outcomes of tissue and microenvironments directly without downstream processing algorithms. By employing weakly supervised learning methods based on tissue-scale labels, InterSTELLAR is designed to simultaneously predict tissue outcomes and detect disease relevant microenvironments. We apply InterSTELLAR to an open-source breast cancer IMC dataset [4] and show that it can accurately characterize patient tissue clinical subtypes. Moreover, by utilizing identified cell communities with high diagnostic value, InterSTELLAR can benefit microenvironment exploration and correlative patient outcomes. We demonstrate the InterSTELLAR workflow using the breast cancer IMC dataset [4], but the method can be easily modified to analyze any other types of highly multiplexed imaging data, such as CODEX, CyCIF and MIBI.

## 2 Results

### 2.1 Interpretable spatial cell learning (InterSTELLAR) principle

Through the modeling of cell spatial interactions of patient tissue specimens, InterSTELLAR aims to predict clinically relevant tissue subtypes and corresponding cell communities (Methods). Here, we focus on a key feature set in breast cancer pathology assessment, including hormone status and growth factor receptor expression, to investigate these cell and cell organizations in aggressive triple negative breast cancer (TNBC). As shown in Fig. 1A, with spatial cell locations, InterSTELLAR first builds undirected graphs per sample to represent topological relationships between cells. In these graphs, a single cell is denoted as a node, and the edges between the cells establish the cell-cell communications. Thereafter, by aggregating marker information (Supplementary Table 1) and interaction between cells, InterSTELLAR builds a four-layer graph convolutional neural network to classify the breast cancer tissues as healthy, TNBC and non-TNBC subtypes (Fig. 1B). In particular, the two graph convolutional layers not only learn from the highly multiplexed cellular data, but also exploit the features from the cell interactions. This strategy enables the identification of spatial domains related to specific clinical subtypes.

**Figure 1.**
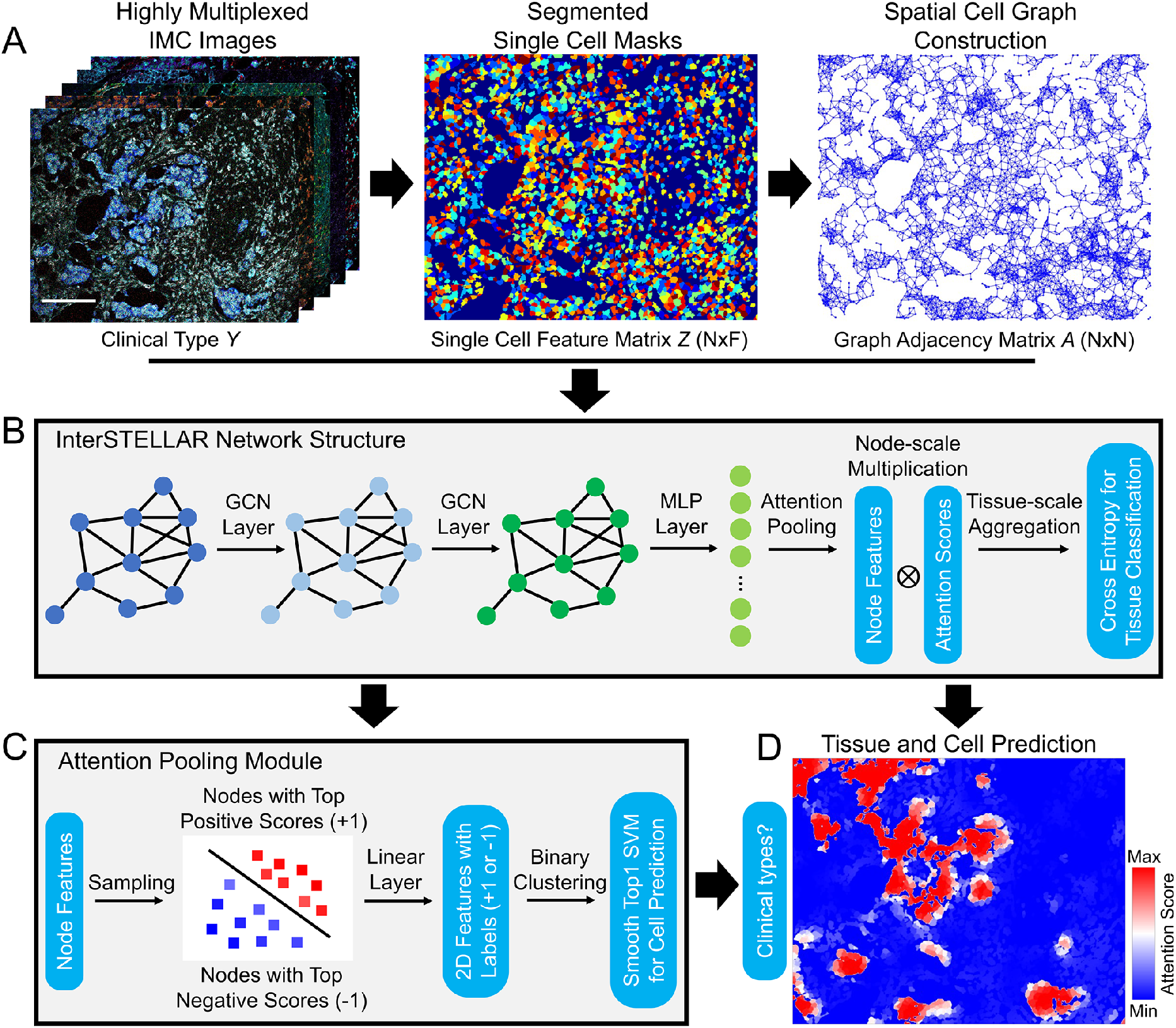
Overview of InterSTELLAR. (A) Highly multiplexed images are segmented, followed by high dimensional single cell data extraction. Simultaneously, undirected graphs are constructed based on the spatial cell locations per tissue. (B) Single cell feature matrix Z, adjacent matrix A and tissue label Y per tissue are fed into InterSTELLAR. As a result, both single cell data and spatial cell interactions are taken into consideration in the training process. (C) In the self-attention pooling module, single cell features with top highest and lowest attention scores are extracted, labeled and trained by a binary classifier, such that the features are linearly separable during inference. (D) A cell-based attention heatmap generated by InterSTELLAR, in which the attention scores per cell are positively correlated with their contribution to the tissue classification results. The rationale comes from the tissue-level aggregation of attention-based pooling, which computes the tissue representation as xthe sum of all cells in the tissue weighted by their respective attention score. Cells from the same neighbourhood share similar attention scores so that functionally they form distinct communities. Scale bar: 172 *μ*m.

To achieve interpretable cell community learning, a self-attention pooling module is embedded between the last hidden layer and the output layer. By adopting a weakly supervised strategy in [21] (Fig. 1C), the network learns to assign attention scores to cells. This attention score quantifies the contribution of cellular spatial groups to the final tissue type predictions (Fig. 1D). This metric of the attention score can be interpreted with the tissue-scale aggregation rule of attention-based pooling, which computes the whole tissue representation as the weighted average of all cells in the tissue by their respective attention score. In this sense, the higher the attention score, the greater the contributions of the corresponding cells to the tissue representation will be, and vice versa. Therefore, the attention scores can quantify the diagnostic value of cell communities to delineate tissue clinical types.

### 2.2 InterSTELLAR achieves accurate clinical type classification and cell-scale characterization

We first evaluated InterSTELLAR on clinical tissue subtype classification. From the data, 73 samples were selected as a test set, and the remaining 293 tissue were trained with a 10-fold cross validation strategy [22]. For reference, InterSTELLAR was benchmarked in comparison with a fully connected neural network (FNN), Random Forest algorithm, and Support Vector Machine (SVM) algorithm (Methods). To account for the label imbalance issue (83 healthy, 49 TNBC and 234 non-TNBC tissues), both balanced accuracy and macro-averaged F1 score were used to evaluate the classification performance (Methods). As demonstrated in Fig. 2A, InterSTELLAR outperforms the three comparator algorithms in both cross validations and testing results. FNN most closely compares to InterSTELLAR of the three; however, FNN lacks information about the relative spatial arrangement of cells within a tissue, so that only the marker features are available in the learning task. Under this condition, some tissues with clinical outcome-relevant spatial structures may be misclassified. As a result, the FNN approach cannot outperform InterSTELLAR. For Random Forest and SVM, performances in cross validation are close to FNN; however, their accuracies decline severely in the independent test set. We infer that the loss of cell-scale information results in the poor performance of generalization of these two algorithms.

**Figure 2.**
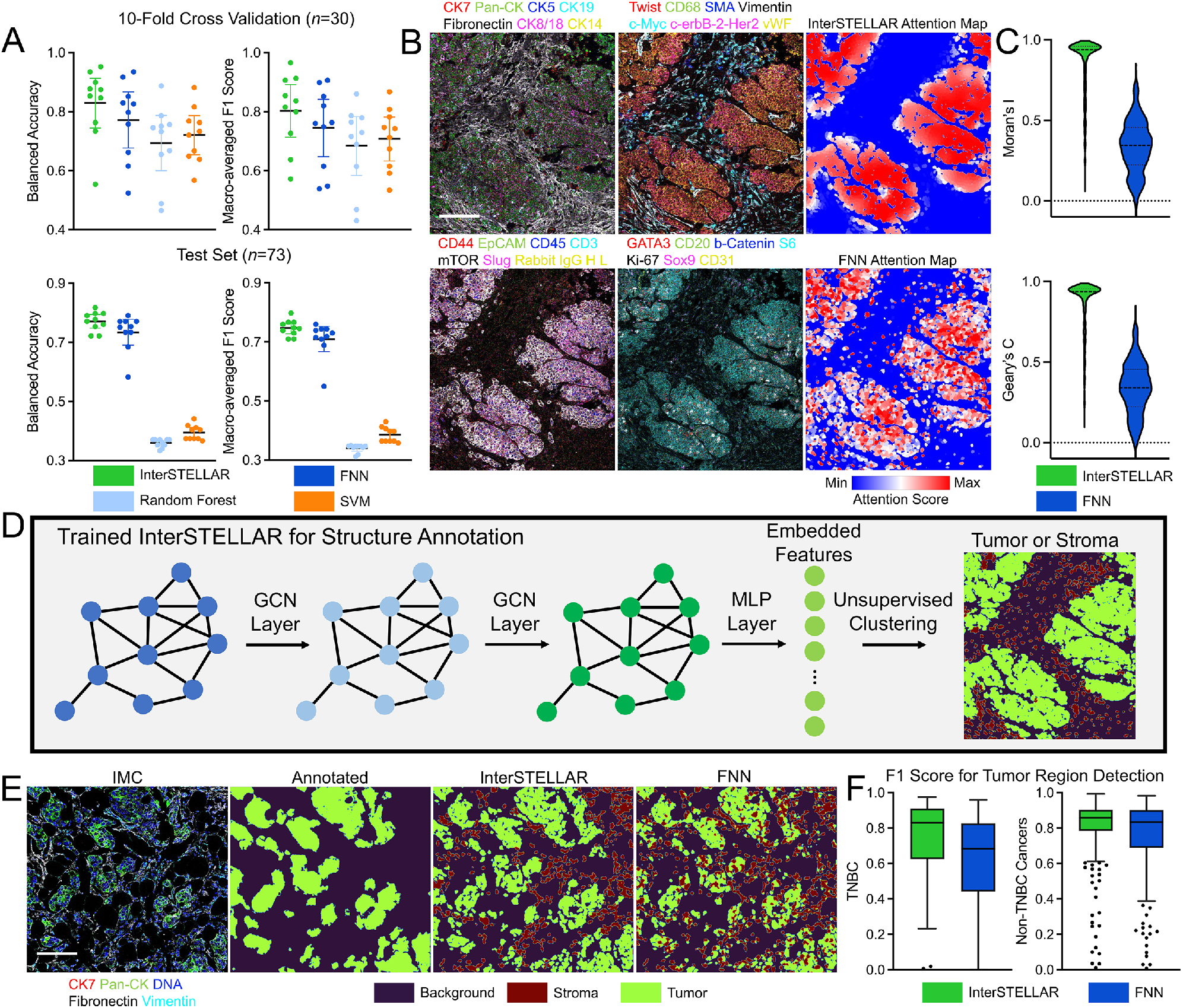
Evaluations of InterSTELLAR. (A) InterSTELLAR is more accurate than FNN, Random Forest and SVM algorithms on tissue clinical type classification, validated by 10-fold cross validation (n = 30 per fold) and an independent test set (n = 73). (B) Highly multiplexed IMC images and the corresponding attention heatmaps generated by InterSTELLAR and FNN. (C) Moran’s I and Geary’s C statistics indicate InterSTELLAR achieves higher spatial correlation than FNN in terms of attention scores. (D) Schematic of generating tumor and stroma masks. Embedded features are extracted from a trained network and then classified by unsupervised clustering algorithms. Here, K-means clustering with K = 2 is utilized. (E) Highly multiplexed IMC images and the corresponding tumor and stroma masks generated by manual annotations, InterSTELLAR and FNN. (F) InterSTELLAR performs better than FNN on tumor region identifications of both TNBC and non-TNBC tissues in terms of F1 score. Scale bar: (B) 145 *μm*; (E) 158 *μm*.

Subsequently, we analyzed the predicted cell-scale attention scores of InterSELLAR, and again benchmarked performance to FNN. Random Forest and SVM are omitted from benchmarking, because they are not feasible methods to predict tissue and cell-scale outcomes simultaneously, due to heterogeneous cell number per field of view in the tissue data set. Both InterSTELLAR and FNN can construct cell-based heatmaps through attention score values (Fig. 2B). However, with more homogeneous distribution of attention scores, InterSTELLAR is superior than FNN on interested community identification. In fact, the attention scores in the same community should be continuous due to the cell-cell communications, such that the cells from the neighbourhood make similar contributions. That is, the attention score of a single cell should be spatial correlated with those of its neighbours. To evaluate the spatial cell correlation per tissue, we benchmarked InterSTELLAR and FNN in terms of Moran’s *I* and Geary’s *C* statistics (Methods). Moran’s I measures how one region is similar to other spots surrounding it. If the regions are attracted by each other, it implies the regions are not independent. Therefore, it is positively correlated with the spatial correlation relationship. Similar to Moran’s I, Geary’s C is also used for the evaluation of spatial autocorrelation. Both the quantitative results of these two metrics in Fig. 2C validate that InterSTELLAR achieves higher spatial correlation, which endorses its improved ability to identify interested communities.

With cell neighbourhood information integrated in training, we hypothesized that the latent embedding features of InterSTELLAR could reflect biologically meaningful information about the tissue structure. Therefore, we proposed to classify all the cells per cancer tissue to either tumor or stromal regions, and then compared the segmentation results with manually labeled tumor region masks by Ilastik [4, 23] (Fig. 2D). Specifically, a K-means clustering algorithm with *K*=2 was utilized to cluster the latent cell embedding features from the last hidden layers of the trained InterSTELLAR network (the input features to the attention pooling module). As a reference, the latent cellular features from the FNN were also clustered with the same approach. Visual inspection and quantitative evaluation confirm InterSTELLAR is superior at capturing tumor regions in both TNBC and non-TNBC tissues (Fig. 2E and F). Indeed, the assembled tumor organizations from InterSTELLAR are much closer to the manual annotations. Interestingly, the FNN has a particular weakness for errantly identifying isolated epithelial cells as tumor cells within the stromal region. Together, these analyses reveal the capabilities of InterSTELLAR over reference methods to identify and interpret class-level features.

### 2.3 Attention Mapping by InterSTELLAR Across Cancer Tissue Types

Pathological categorization of breast cancer is critical for patient care and is typically accomplished with well-established markers for hormone status, HER2 expression and tissue morphology. We next investigated whether InterSTELLAR enables microenvironment characterization for pathologically distinct clinical types of breast cancer from the same IMC dataset. We defined high- and low-attention regions by segmenting the attention heatmaps with the median attention score of each sample (Fig. 3A). With these binary masks, we calculated the percentages of immune, stromal and epithelial cells in high- and-low attention regions for healthy, TNBC, and non-TNBC patient tissues. The attention regions for each tissue type show distinguishing compositions for these cell types (Fig. 3B). First, we observed that healthy tissues have higher stromal cell proportion in both attention regions than any cancer tissues. Importantly, high-attention regions within cancer tissues contain more epithelial cells than of all the remaining regions. Comparatively, the low-attention regions correspond with an increased stromal cell presence than the high-attention regions from the same tissues. Low-attention regions from all three clinical types contain similar portions of epithelial cells. Interestingly, TNBC tissues had the highest proportion of immune cells compared to healthy or non-TNBC tissues, revealing higher immune activity in the TNBC tissue microenvironment. In non-TNBC specimens, low-attention regions have a higher proportion of immune cells than healthy tissues, but there are no appreciable differences in immune cell percentages of the high-attention regions. In sum, the variations of cell composition by attention scores can distinguish the microenvironments from different clinical subtypes of breast cancer.

**Figure 3.**
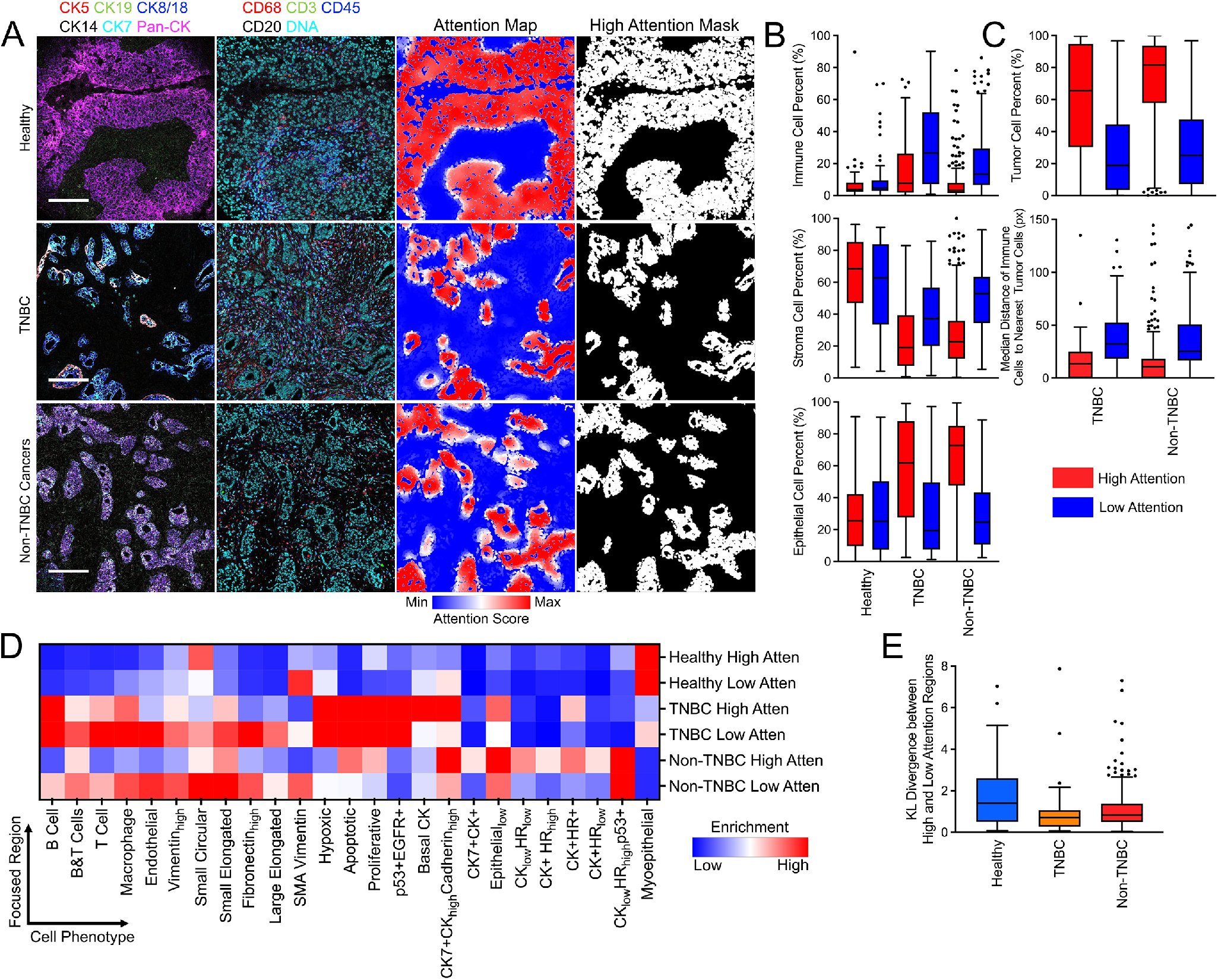
InterSTELLAR characterizes the breast cancer tissue microenvironments from different clinical subtypes. (A) Highly multiplexed IMC images of healthy, TNBC and non-TNBC tissues as well as their corresponding attention heatmaps and segmented high attention region masks. (B) The percentages of immune, stroma and epithelial cells in high and low attention regions from healthy, TNBC and non-TNBC tissues. (C) The percentages of tumor cells and median distance of immune cells to their nearest tumor cells in high and low attention regions from cancer tissues. (D) Mean cell density per phenotype from high and low attention regions of all the tissues. (E) KL divergence between high and low attention regions regarding distributions of cell density per phenotype. Scale bars in (A): Healthy: 195 *μm*; TNBC: 175 *μm*; Non-TNBC cancers: 183 *μm*.

Next, we calculated the percent of tumor cells in high- and low-attention regions from cancer tissue specimens (Fig. 3C). As expected, high-attention regions are occupied by a greater number of tumor cells than low-attention ones. As an example, previous reports have suggested that tumor-infiltrating lymphocytes may be an indication of TNBC [24]. Additionally, high-attention regions have higher ratios of tumor cells in non-TNBC tissues than TNBC, which is inversely related to the immune cell ratios for these tissues. This suggests more frequent immune-tumor cell interactions in TNBC tissues. Moreover, we calculated the median distance of immune cells from both high- and low-attention regions to their nearest tumor cells (Fig. 3C). The results reveal that immune cells in high-attention regions are in closer proximity to tumor cells. Therefore, more frequent immune-tumor interactions, such as tumor-infiltrating lymphocytes, are expected in these regions.

### 2.4 InterSTELLAR captures tissue microenvironmental features from different clinical subtypes

Different clinical types of breast tissue are characterized by distinctive cell phenotypes compositions, for example, myoepithelial cells in healthy tissues and extensive proliferative cells in TNBC. Now, we ask whether distinct attention regions can also be characterized by distinct cell-type compositions, even in the same tissue. We adopted a cell phenotype approach for breast cancer tissues as previously described [4]. To conduct more detailed phenotype analysis, we calculated the cell density per phenotype for all the attention regions (Supplementary Fig. 1). The normalized distributions of the mean cell density of each phenotype are summarized in Fig. 3D. This demonstrates that in TNBC high-attention regions there are more Basal CK and Epithelial_low_ cells, but fewer myoepithelial cells, compared to TNBC low-attention counterparts. To quantify such differences, we calculated the Kullback–Leibler (KL) divergence of the phenotype distributions between high- and low-attention regions per sample (Fig. 3E). The results reveal that almost all the KL divergence values are away from 0, indicating that cell phenotype compositions enable expected attention score differentiation. Interestingly, more diverse distributions are noticed in healthy tissues rather than cancer types. We inferred that this reveals more heterogeneity in the microenvironments in healthy tissues.

### 2.5 InterSTELLAR uncovers single-cell pathology groups associated with patient survival

Beyond the microenvironment and spatial characterization for various clinical types, we were curious whether InterSTELLAR could benefit single-cell pathology (SCP) analysis. Using the unsupervised Phenograph algorithm [25, 26], we grouped patient cancer tissues on the basis of cell phenotype densities within high-attention regions and identified 8 SCP subgroups (Methods), which are named according to their dominant phenotypes (Fig. 4A and Supplementary Fig. 2). The dimensions of the density data were also reduced by uniform manifold approximation and projection (UMAP) algorithm for visualization (Methods). Through inspection, the tissues are clustered by their distinct phenotype compositions (Fig. 4B).

**Figure 4.**
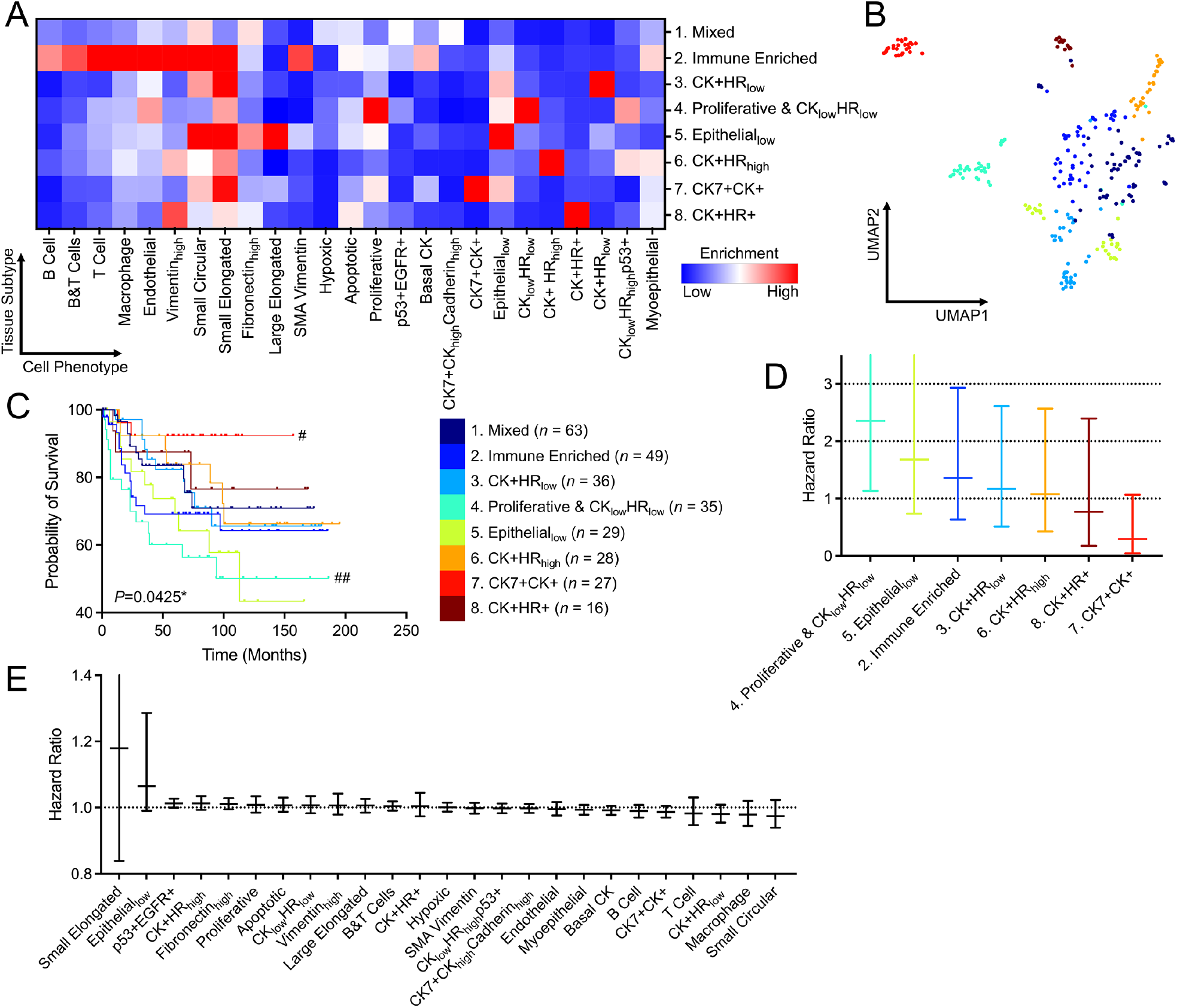
InterSTELLAR characterizes distinct clinical outcomes for SCP subgroups. (A) Mean cell density per phenotype of high attention regions from various SCP subgroups. (B) UMAP plot of the tissues labeled with their corresponding SCP indexes. (C) Kaplan-Meier curves of overall survival for each subgroup (n = 283) on the basis of cell density per phenotype of high attention regions, with \**P*<0.05. #*P*<0.05, ##*P*<0.01 represent the statistical significance of a single subgroup compared to all other samples. (D) F values for overall survival analysis from different clustering strategies. In (C, D), *P* values were calculated through two-sided log-rank test. (E, F) Relative hazard ratios and 95% confidence intervals of disease-specific overall survival for cell densities per phenotype and SCP subgroups estimated using a Cox proportional hazards model. Reference group 1: Mixed for SCP subgroups.

Importantly, the presence of these subgroups have significantly different clinical outcomes in overall survival (Fig. 4C), validated by a log-rank test with *P*=0.0425 on Kaplan-Meier curves (Methods). In particular, the CK7+CK+ subset in high attention regions defined patients with favorable clinical outcome, while the presence of proliferative CK_low_HR_low_ was associated with adverse overall survival. Hormone receptor and HER2 subtypes and tumor grade were associated with prognosis, as expected (Supplementary Fig. 3). Most remarkably, SCP features identified cohorts that were independent from clinical subtype or tumor grade (Supplementary Fig. 3A) with distinct survival results (Supplementary Fig. 3B). Also compared to clinical subtype and tumor grade results, SCP subgroup analysis with InterSTELLAR allows a higher-resolution tissue characterization paradigm (4 and 3 vs 8).

Based on the observation that SCP analysis within high-attention regions appeared associated with clinically significant outcomes, we examined whether this association was also present for low-attention regions or whole tissue analysis. Applying the approach above for low-attention regions (*P* = 0.8424) or whole regions (P=0.1381), there was no statistically significant difference in assessing outcome (Supplementary Fig. 4 and Supplementary Table 2). We conclude that high-attention regions have greater diagnostic value such that they are more relevant to contribute to patient outcomes. Since high-attention regions contain a higher proportions of epithelial cells, we used the same unsupervised clustering approach (Methods) based on only epithelial cells from high-attention regions, or clustering using epithelial cells from the entire tissue sample (Supplementary Fig. 4 and Supplementary Table 2). Both of these approaches based on using only epithelial cells did not reach the significance of all cell types in the high-attention region, indicating that indeed the spatial organization of multiple cell types are essential in microenvironment analysis and interpretation.

Inspecting the subgroups identified through SCP analysis of high-attention regions, prognostic groups become apparent (Supplementary Fig. 5). SCP Group 4 Proliferative & CK_low_HR_low_ has very poor prognosis with less than 70 percent overall survival at 5 years (Cox proportional hazard HR=2.36, CI: 1.13–4.99, Fig. 4D), while SCP Group 5 Epithelial_low_ (HR=1.68, CI: 0.74–3.76) and Group 2 Immune Enriched (HR=1.36, CI: 0.63–2.93) are moderately unfavorable. In contrast, SCP Group 7 CK7+CK+ exceeds 90 percent overall survival beyond 10 years (HR=0.29, CI: 0.05–1.07). SCP Groups 1, 3, 6 and 8 have intermediate risk. Alternatively, when analyzing each single cell phenotype within the high-attention region but without clustering by tissue composition, no individual cell type was directly correlated with outcomes (Fig. 4E). Thus, SCP subgroups clustered by the tissue composition from high-attention regions provide an innovative approach to inform prognosis.

## 3 Discussion

With the expansion of novel multiplexed technologies for the characterization of cellular context in health and disease, graph-based deep learning algorithms have begun to be investigated on high dimensional single cell data. In this work we present InterSTELLAR, a GNN framework that can predict patient tissue outcomes and disease relevant communities simultaneously. We introduce and evaluate InterSTELLAR using an open-source breast cancer IMC dataset. Cell microenvironments per tissue are first modeled as graphs, and the nodes represent cells and the edges represent inter-cell communications. Then, graph convolutional layers are applied to extract cell interaction features, followed by a self-attention pooling module to learn the cell-based contribution to clinical outcomes. InterSTELLAR achieves higher accuracy than traditional machine learning algorithms, including Random Forest, SVM, and a FNN framework which neglects spatial cell information. Moreover, InterSTELLAR performs better than FNN in terms of interested community identification. We infer that modeling cell communications is crucial to accurate tissue characterization.

We performed validation studies indicating that InterSTELLAR can capture distinct tissue microenvironment features from healthy and breast cancer tissues. With high- and low-attention regions segmented by median attention score values, we observed disease-specific composition of immune, stroma and epithelial cells per tissue. This approach removes manual segmentation of regions of interest by an expert pathological reader to identify regions within a field of view of high (and low) value for correlating with microenvironmental subclasses. Our analyses revealed that greater numbers of tumor cells are localized in high-attention regions of cancer tissues, which potentially distinguish communities by diagnostic value. Propinquity of immune and tumor cells from high-attention regions is indicative of active immune-tumor interactions in these domains. Furthermore, the heterogeneity of low- and high-attention regions based on cell phenotypes reveals the ability of InterSTELLAR to discriminate clinically important aspects of the tumor microenvironments.

Through SCP subgroup analysis, we find that InterSTELLAR can establish a mapping relationship between tissue microenvironment and breast cancer patient prognosis. Specifically, each subgroup has unique cell community features in high attention regions, which are significantly associated with the survival outcomes. By contrast, other attention regions or areas of epithelial cells do not have this association. Moreover, both Kaplan-Meier curves and Cox proportional hazards modeling suggest distinct survival outcomes of the subgroups.

The results discussed in this work demonstrate the value of InterSTELLAR for high-dimensional spatial cell data to investigate cellular communities of interest. A particular strength of the INTERSTELLAR approach is the flexibility to apply this method across multiple platforms for highly multiplexed imaging, because it is capable of extracting cell marker features and tissue graphs independent of the imaging modality. As a supervised learning framework, InterSTELLAR requires substantial training data to guarantee the generalization of the trained model and careful minimization of batch effects on marker staining to avoid overfitting and to prevent degradation of prediction performance. The current study analyzes a large, recent cohort of patient tissue data, but still remains limited in providing sufficient size for validation of the intriguing findings suggested by SCP analysis. Future advances in imaging platforms and staining methods that improve the ability to generate affordable, very large data sets will be an exciting opportunity for wider implementation of the InterSTELLAR approach. The proposed framework can be applied to other highly multiplexed imaging techniques and diseases for enhanced downstream analysis. To conclude, InterSTELLAR is a versatile GNN framework for highly multiplexed imaging data that simultaneously classifies tissue types by clinical classes and predicts disease-relevant cell communities. Most importantly, by exploiting cell communities with high diagnostic values, it enhances the characterization of patient tissue microenvironments.

## 4 Methods

### 4.1 Dataset description and pre-processing

We applied the InterSTELLAR framework on an open-source breast cancer IMC dataset [4]. The dataset consists of 381 tissues with 35 cell markers, as well as segmentation masks, single cell data, cell phenotypes and tumor-stroma masks as well as tissue clinical subtype and patient survival information. To focus on the effects of antibody markers and to increase robustness, DNA markers and tissues with too few cells (less than 50) were removed, leaving 366 tissues with 30 markers (Supplementary Table 1). Specifically, there are 83 healthy, 49 TNBC and 234 non-TNBC tissues.

For a specific cell marker *c*, denote the single cell data 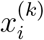 as the mean expression value of each cell *i*, and the array of all expression values 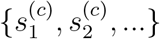 as *S*^(*c*)^. We then normalized the single cell data using log transformation:

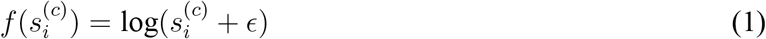

where *ϵ* represents a small value. Here, we set it as 10^-4^. Next, we calculated the z-score of the normalized expression value:

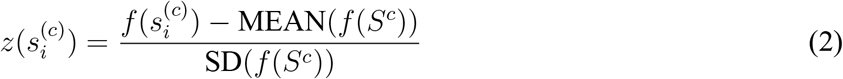

After normalization per marker, for each tissue we obtained a cell-by-marker expression matrix *Z^N×F^* filled with pre-processed expression values 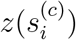, in which *N* is the total cell number for the tissue, and *F* is the marker number.

### 4.2 Graph construction

InterSTELLAR is built upon undirected graphs. To construct graphs from tissues, the set of cells are represented by a set of discrete points located at cellular centroids. The 2D coordinates of these cellular centroids (*x, y*) are determined by the segmentation masks of the corresponding cells. Then, we regard each tissue as a single graph, in which each cell is a node of the graph. The node features are the matrix of marker expressions *Z^N×F^*. The edge between any two nodes determines whether the nodes are connected. The Euclidean distance between any two nodes *u* and *v* is calculated as:

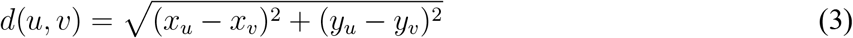

Considering any two cells in a tissue, the longer their distance, the less their inter-communications will be. We define the weight of each edge (*u, v*), which is negatively associated with their distance *d*(*u, v*):

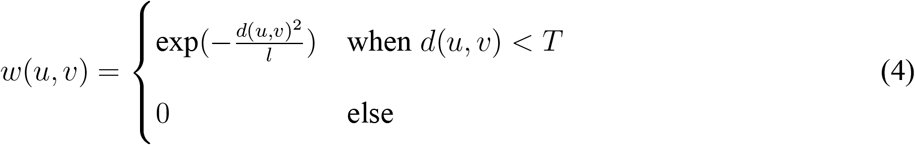

In Eq. (4), the hyperparameter *l* and *T* determines how rapidly the weight decays as a function of distance. Here, we set *T* as 40 *μm.* This is approximately twice the size of a regular cell as it assumes that cells have to be within reach to each other to interact [14]. *l* is empirically set as 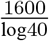 so that *w*(*u, v*) approaches 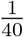 when *d*(*u, v*) approaches the threshold. Therefore, the graph adjacency matrix *A^N×N^* with a shape of *N* × *N* is built, in which *w*(*u, v*) is the matrix element. Finally, there are a node feature matrix *Z^N×F^*, a graph adjacency matrix *A^N×N^* and a clinical type label *Y* ∈ {Healthy, TNBC, Non-TNBC cancers} per graph as inputs to InterSTELLAR.

### 4.3 Network structure

For each constructed graph, a collection of (*Z^N×F^, A^N×N^, Y*) are fed into InterSTELLAR. The inputs *Z^N×F^* and *A^N×N^* first pass through two graph convolutional modules, and the node feature matrix is transformed as 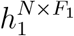 and 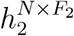, respectively. Each module consists of a graph convolutional layer [27], a layer normalization module [28] and a scaled exponential linear unit (SELU) [29]. Subsequently, 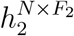 is fed into a fully connected layer followed by another SELU module, and the output is 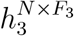.

To achieve tissue-scale classification and cell-scale interpretable learning, a self-attention pooling module, modified from [21], is embedded between 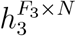 (the transpose of 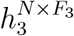) and the final output. The attention score of the *i*-th cell is defined as Eq. (5). As a result, the tissue-scale representation aggregated per the attention score distribution is defined as Eq. (6), in which ⊗ denotes element-wise multiplication.

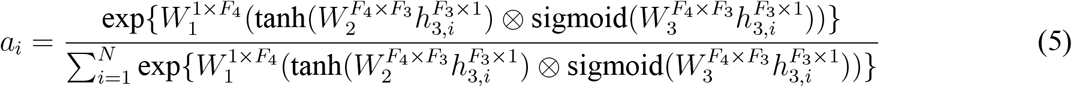

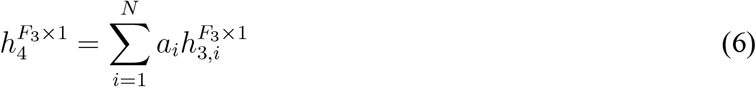

The attention score *a_i_* is the cell-scale contribution to the final tissue-scale output. Therefore, *a_i_* is positively correlated with the diagnostic value per cell. Finally, 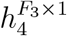 is further transformed as 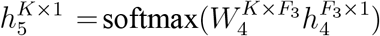, and cross entropy is set as the loss function 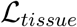 between the tissue-scale prediction and the label *Y*, in which *K* is the tissue class number.

In the attention pooling module, an additional binary clustering objective is introduced so that class-specific features can be learnt [21]. During training, a collection of 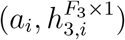 is sorted according to the value of *a_i_*, in which *i* = 1,2,…, *N*. Then, the pairs of 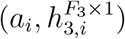 with *Q* top highest and lowest *a_i_* are selected. Next, *K* separate fully connected layers are utilized to process the selected 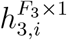 for each class, respectively (*k* = 1,2,…, *K*):

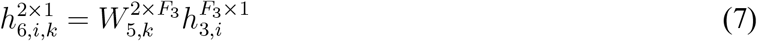

Regarding the training of each classifier, the 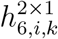 with the *Q* top highest *a_i_* are attached with positive labels (+1) while those with the *Q* top lowest α*i* are attached with negative labels (−1). The smooth top-1 SVM loss is selected as the loss function 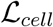 for this cell-scale learning task, because it has been empirically shown to reduce over-fitting under the conditions of noisy data labels or limited data [30]. Note that the labels are independently generated in each iteration. Intuitively, the sub-training task in each of the *K* classes is supervised by the corresponding tissue-scale label. Consequently, the cell communities with high attention scores are expected to be positive evidence for its current tissue label; By contrast, the communities with low attention scores are the negative evidence. Therefore, this sub-training task can be regarded as a constraint for the cell-scale feature 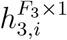, such that the features favoring the correct outcome are linearly separable from those uncorrelated ones. The overall loss function 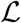, as Eq. (8) shows, is the weighted sum of 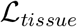 and 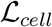, in which *η* ∈ [0,1] is the tissue-scale weight.

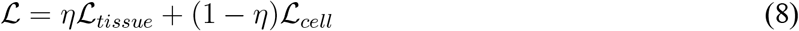

### 4.4 Model training

We preset that *F* = 30 and *K* = 3 for the breast cancer IMC dataset training task. Other hyper-parameters were set as *F*_1_ = 40, *F*_2_ = 40, *F*_3_ = 20, *F*_4_ = 10 and *Q* = 8. The optimal η was found as approximately 0.85 so that the highest accuracy is achieved (Supplementary Fig. 6).

The network was trained using Pytorch [31] (version 1.10.2) and Pytorch-Geometric [32] (version 2.0.4) on a single NVIDIA Quadro RTX 6000 GPU with 24 GB of VRAM. To mitigate the class imbalance in the training set, the sampling probability of each tissue was inversely proportional to the frequency of its label. Of the 366 tissues, 73 tissues were selected as test set, the remaining tissues were trained with a 10-fold cross validation strategy. During the training, the model parameters were updated via Adam optimizer with a learning rate of 3 × 10^-4^ and *l*1 weight decay of 3 × 10^-5^. All the other parameters of the Adam optimizer were utilized with default values. A step learning rate strategy was also applied so that the learning rate was multiplied by 0.9 after each 5 epoches. The network was trained with 30 epoches with a batch size of 8. Each fold took approximately 166 seconds for training. The trained model with the highest validation accuracy was saved for each fold.

### 4.5 Baseline methods

Baseline methods in this work include a fully connected neural network (FNN), a Random Forest classifier and a Support Vector Machine (SVM) classifier. The FNN method is similar to InterSTELLAR – both tissue and cell-scale predictions can be conducted. However, the two graph convolutional layers are replaced by two fully connected layers. In this case, the spatial locations of cells are not taken into consideration so that the adjacent matrix *A^N×N^* is neglected. Thus, FNN can only utilize the single cell data without the information of spatial cell interactions.

Random forest and SVM algorithms are conducted on the basis of composition vector inputs, which are the cell densities of each phenotype per tissue. There are 25 cell phenotypes of the breast cancer IMC dataset (Supplementary Fig. 1). As a consequence, the input per tissue is a 25 ×1 vector. Before training and inference, the inputs are z-score normalized. Compared to InterSTELLAR and FNN methods, only tissue-scale classification is available for Random Forest and SVM, due to the loss of cell-scale information. Note that both random forest and SVM were implemented using scikit-learn [33] (version 1.0.2) with default settings.

### 4.6 Single-cell pathology patient grouping

Patient cancer tissues were grouped on the basis of the cell densities of all the phenotypes in high attention regions using Phenograph [25] with Leiden algorithm [26]. These algorithms were implemented by Scanpy package [34] (version 1.8.2) with 20 nearest neighbours and resolution of 1. All the other parameters were utilized with default settings. These parameters were chosen such that groups of patients from distinct cell type compositions can be successfully separated without limiting statistical power for group comparisons. Similar unsupervised clusterings were also conducted on the basis of the cell densities of all the phenotypes in low attention and all regions, and epithelial cell densities from high attention and all regions. The parameters of Phenograph kept unchanged. The random seeds for the individual runs were recorded.

### 4.7 Uniform manifold approximation and projection (UMAP)

For visualization, high-dimensional cell density data per tissue was reduced to two dimensions using the non-linear dimensionality reduction (UMAP) algorithm [35]. This algorithm was implemented by the umap package (version 0.5.2) after all the inputs were z-score normalized. All the parameters were utilized with default settings. The random seeds for the individual run was recorded.

### 4.8 Survival curves and Cox proportional hazard regression models

Kaplan-Meier survival curves and Cox proportional hazards survival regression models were generated using Prism 9 (GraphPad Software Inc.). The overall survival of patients in different single-cell-defined subgroups was analyzed. Both log-rank tests and Cox proportional hazards models were utilized to investigate the statistical significance of the patient subgroup classification.

### 4.9 Accuracy metrics

To access the tissue-scale classification performance considering the label imbalance effect, balanced accuracy and macro-averaged F1 score were implemented in Fig. 2A with scikit-learn package [33] (version 1.0.2). Segmentation performance for tumor regions in Fig. 2F was accessed by F1 score with scikit-learn package.

Because InterSTELLAR aims to detect the cell communities with high diagnostic values, the predicted attention scores should be continuous through the cell neighbourhood, such that adjacent cells from same communities make similar contributions to the final output. In another word, the attention scores of the adjacent cells should be spatial correlated. We evaluated the correlation by Moran’s *I* [36] and Geary’s C statistics. Moran’s *I* metric is a correlation coefficient that measures how one spot is similar to other spots surrounding it. Its value ranges from −1 to 1. The higher the value, the higher the spatial correlation relationship will be. For the given attention scores, we define the Moran’s *I* using the following formula,

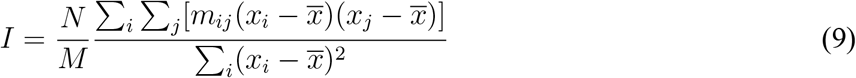

where *x_i_* and *x_j_* are the autention scores of cells *i* and *j*, 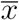 is the mean attention score of a tissue, *m_ij_* is spatial weight between cells *i* and *j* calculated using the 2D spatial coordinates of the spots, and *M* is the sum of *m_ij_*. For each cell, 4 nearest neighbours are selected based on the Euclidean distance between cells. If cell *j* is the nearest neighbour of cell *i, m_ij_* is assigned as 1; otherwise, *m_ij_* = 0.

Geary’s *C* can be also used for the spatial autocorrelation evaluation, which is calculated as

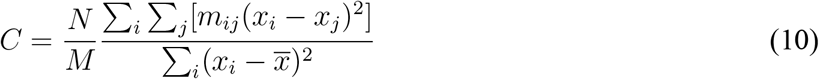

The value of Geary’s *C* ranges from 0 to 2. We transform it as *C*\* = 1 – *C* so that its range will be [–1,1] [17]. Similar to Moran’s *I*, the higher the value of *C*\*, the higher the spatial correlation relationship will be between cells in the same neighbourhood.

### 4.10 Statistical analysis

Other than specially stated, quantitative data are presented as box-and-whisker plots (center line, median; limits, 75% and 25%; whiskers, maximum and minimum). The two-sided log-rank tests were implemented with Prism 9 (GraphPad Software Inc.). Statistical significance at P<0.05, 0.01 are donated by * and ** (Fig. 4C and Supplementary Figs. 4,5) or # and ## (Fig. 4C), respectively.

## Data Availability

We applied InterSTELLAR on the breast cancer IMC dataset [4], which is publicly available. The data links are provided in the corresponding paper. Besides, the pre-processed data is available at https://doi.org/10.5281/zenodo.7527814.

## Code Availability

The InterSTELLAR code used in this study and the corresponding trained weights are publicly available at https://github.com/PENGLU-WashU/InterSTELLAR.

## Acknowledgement

Research support includes NCI K12 CA167540 (K.A.O.) and American Society of Hematology Scholar Award (K.A.O.).

## Supplementary Figures

**Supplementary Figure 1.**
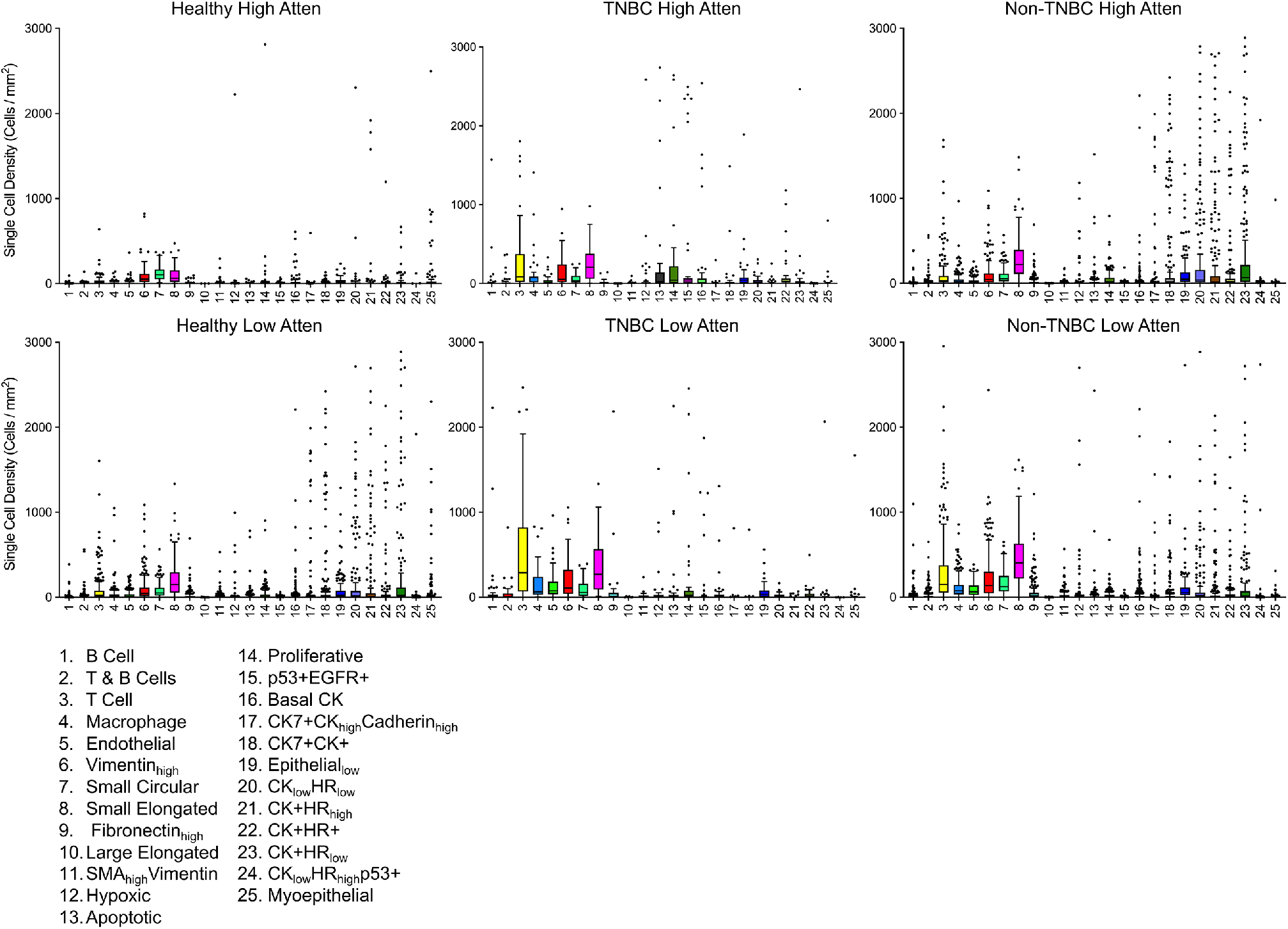
Box plots of cell density per phenotype in high- and low-attention regions from healthy, TNBC and non-TNBC cancer tissues.

**Supplementary Figure 2.**
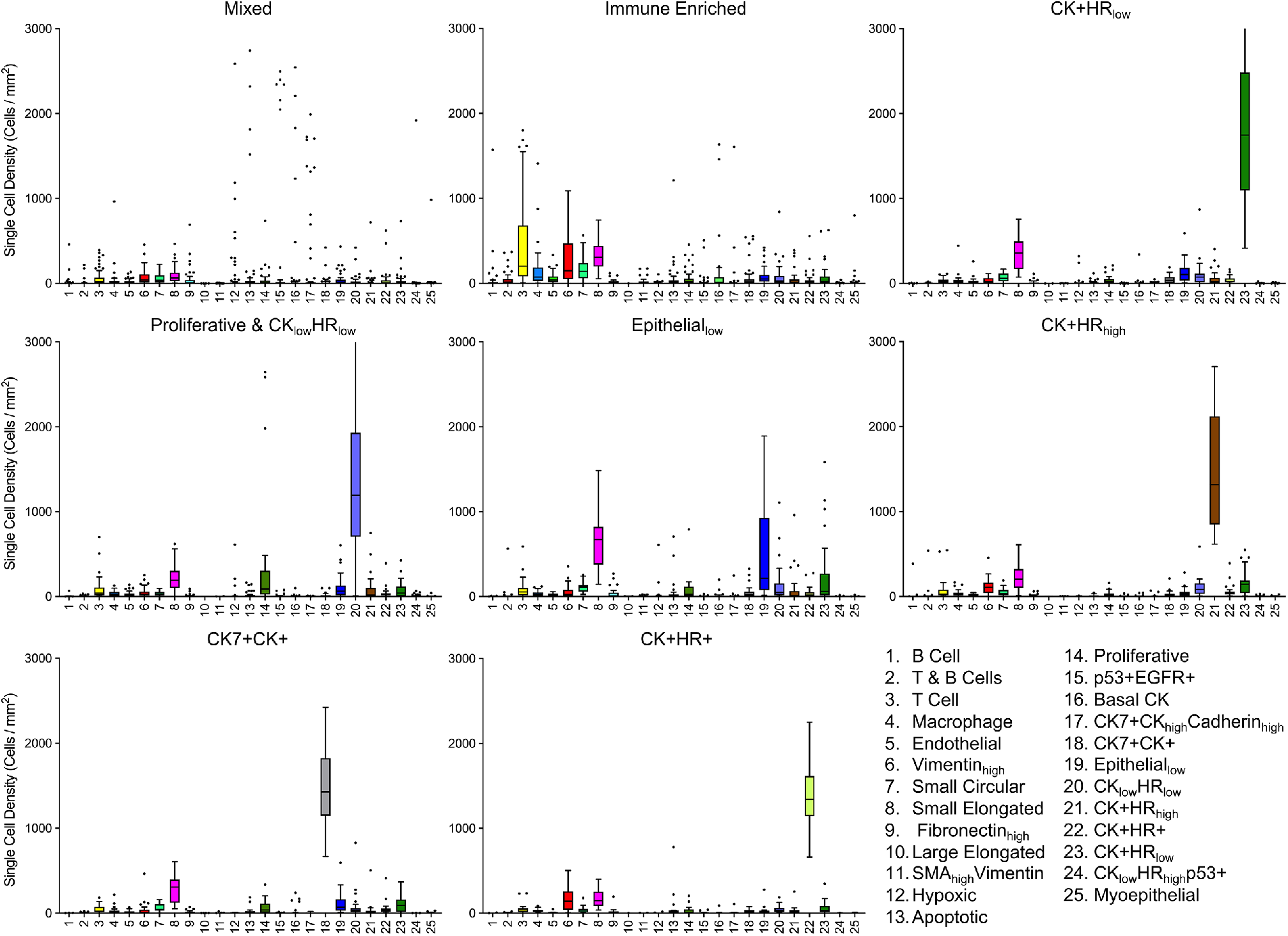
Box plots of cell density per phenotype in high-attention regions from each single-cell patient subgroup.

**Supplementary Figure 3.**
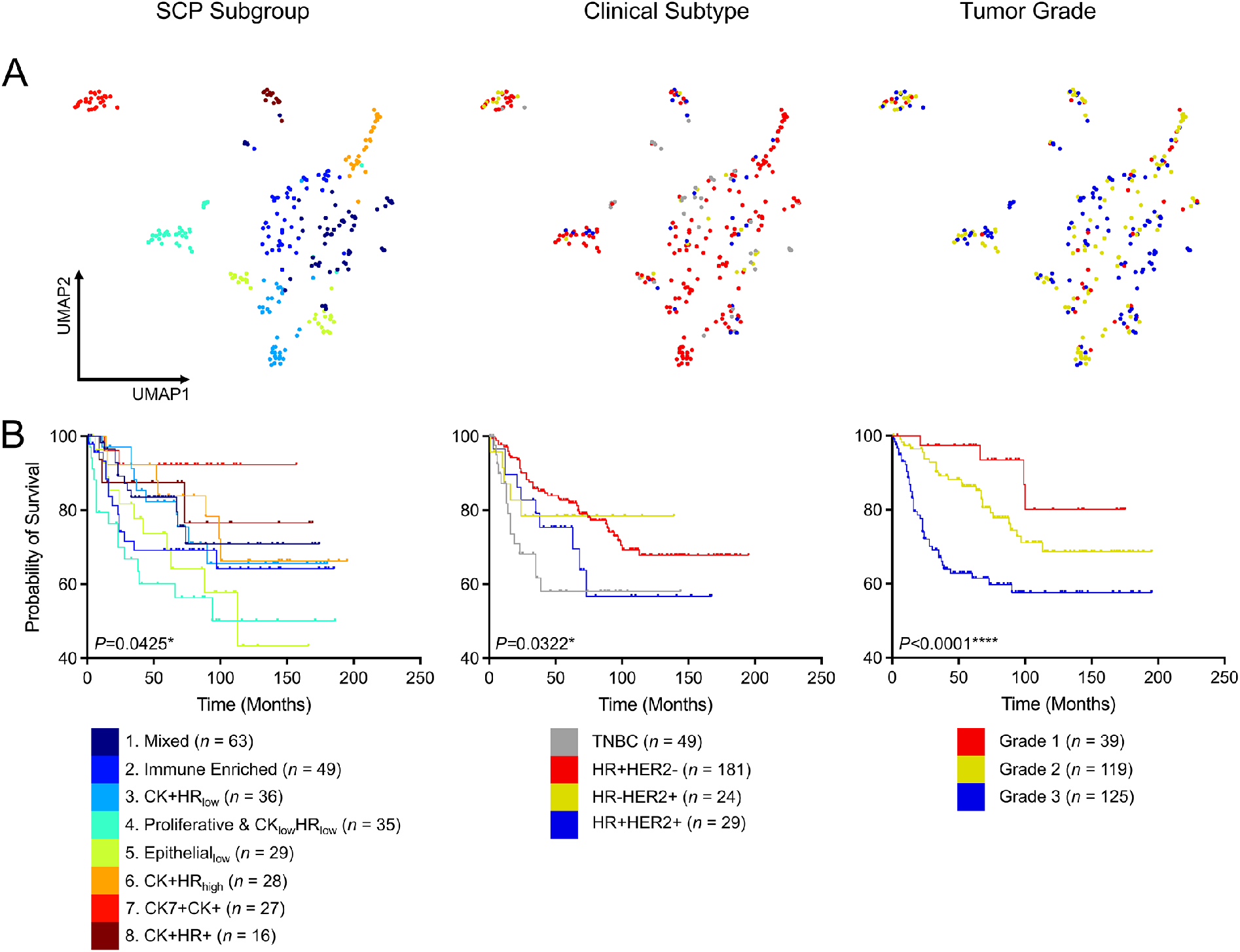
Comparisons of the patient group classified by SCP subgroup, clinical subtype and tumor grade. (A) UMAP plots of the tissues labeled with their corresponding SCP indexes, clinical subtypes and tumor grades. (B) Kaplan-Meier curves of overall survival for each patient group on the basis of SCP subgroup, clinical subtype and tumor grade.

**Supplementary Figure 4.**
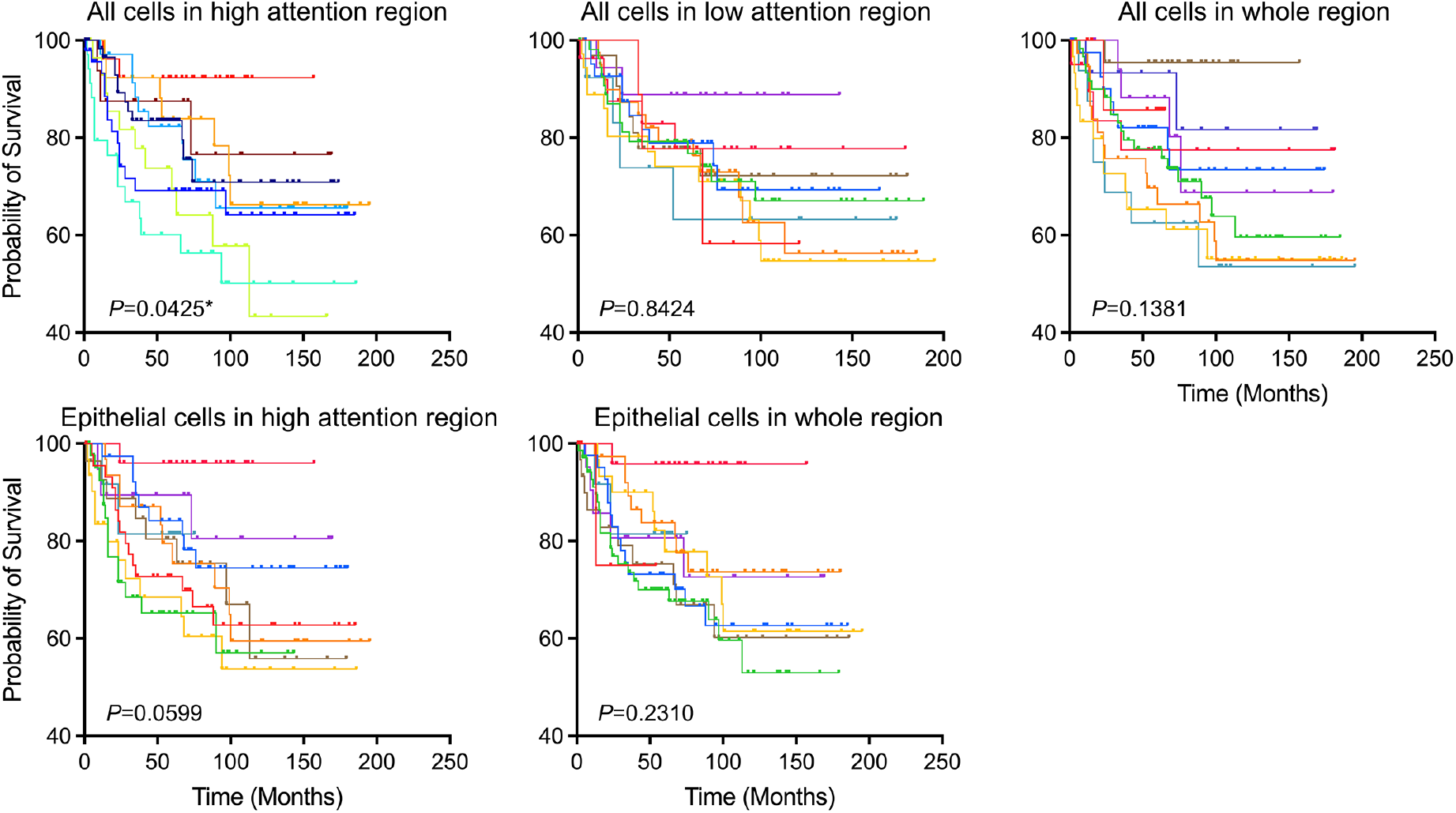
Kaplan-Meier curves of overall survival from different clustering strategies and their corresponding *P* values of log-rank tests.

**Supplementary Figure 5.**
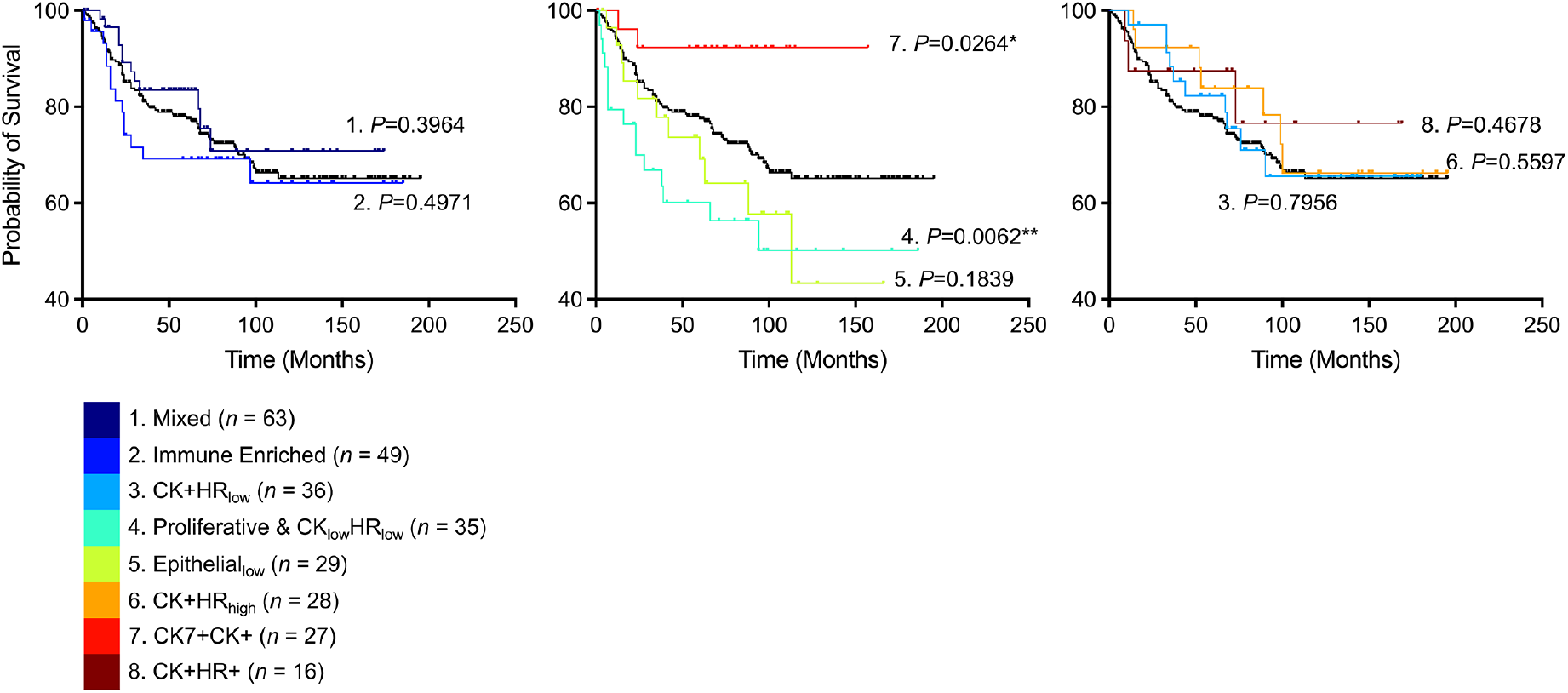
Kaplan-Meier curves of overall survival of each subgroup on the basis of cell density per phenotype of high attention regions. The black curve represents the survival curve of all samples. Each *P* value represents the statistical significance of a single subgroup compared to all the other samples. All the *P* values were calculated through two-sided log-rank tests.

**Supplementary Figure 6.**
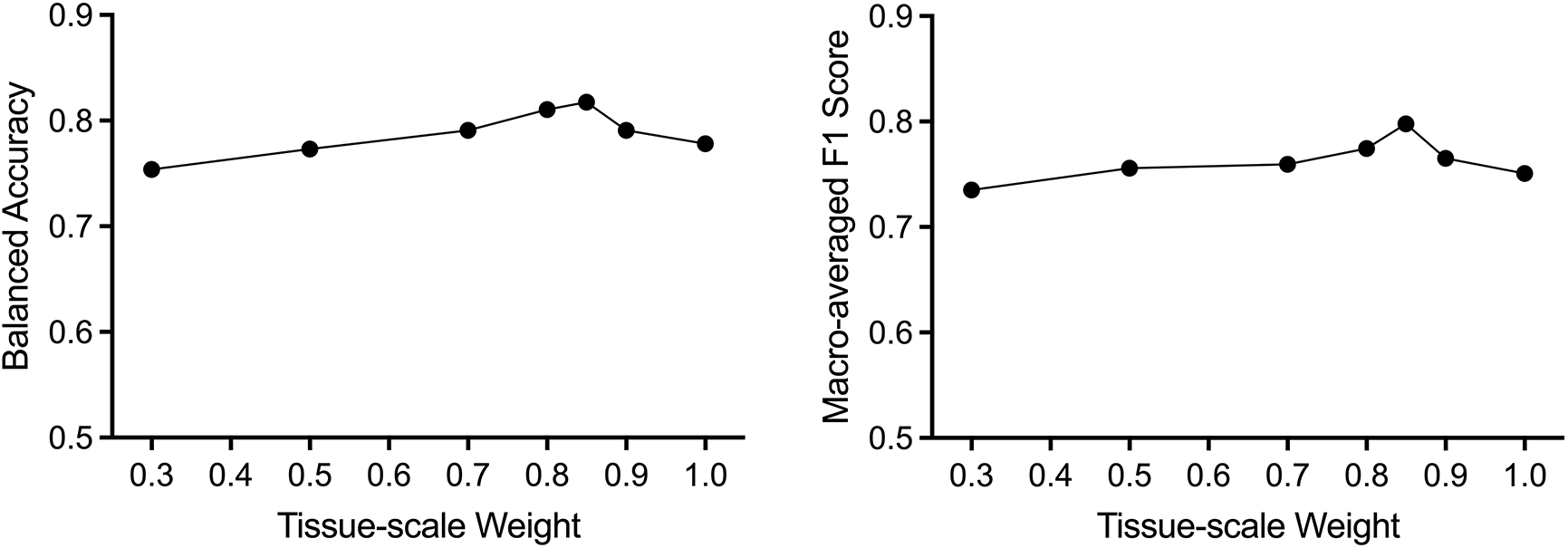
The relationship between tissue-scale weight *η* and clinical type classification performance.

## Supplementary Tables

**Supplementary Table 1.**
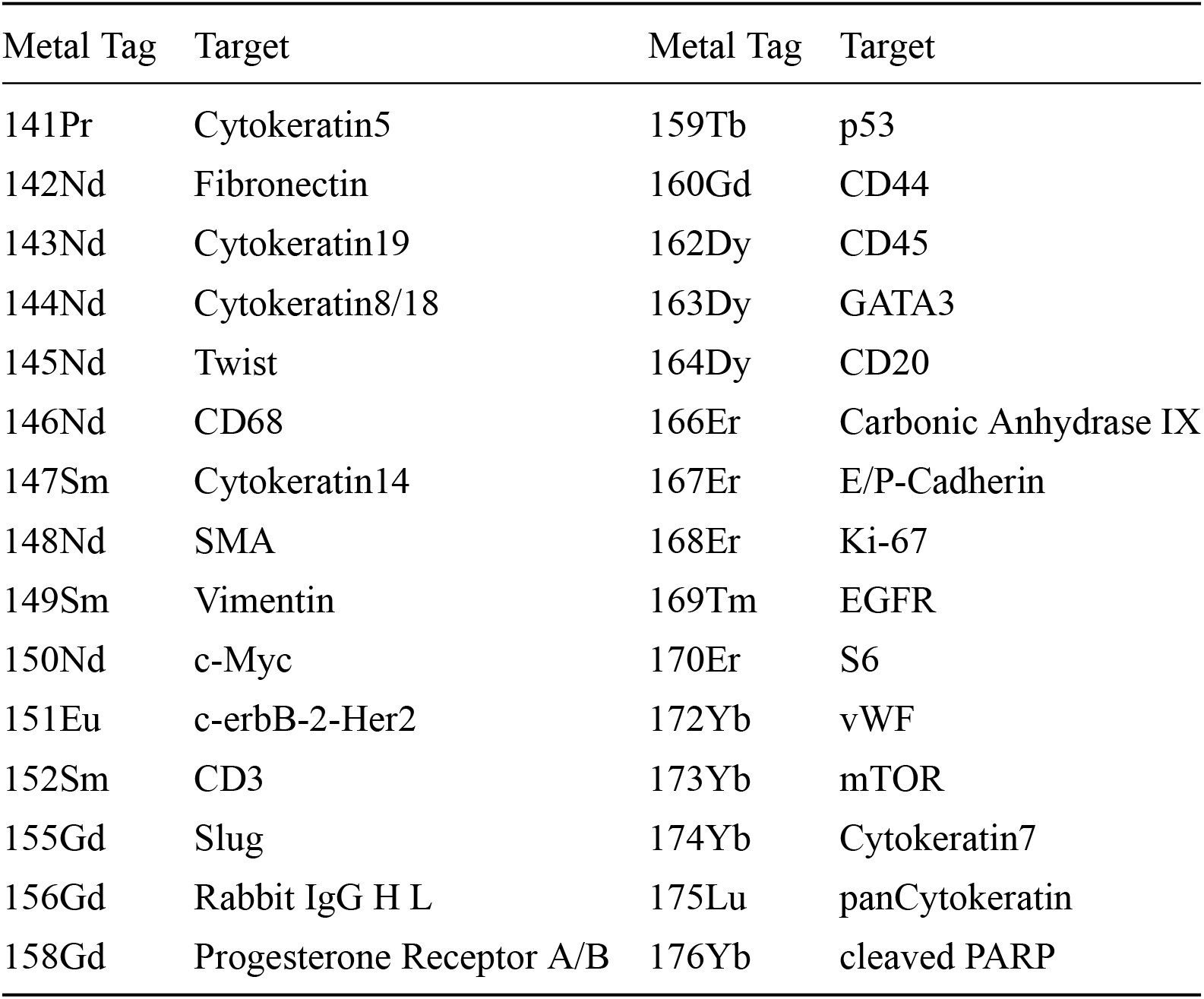
Cell markers used in training and downstream analysis.

**Supplementary Table 2.** *P* values for overall survival analysis from different clustering strategies.

| Cell Types | Region | *P* Value |
| --- | --- | --- |
| All | High attention | 0.0425* |
| All | Low attention | 0.8424 |
| All | Whole | 0.1381 |
| Epithelial cells | High attention | 0.0599 |
| Epithelial cells | Whole | 0.2310 |

## Notes

### Competing Interest Statement

The authors have declared no competing interest.

## References

[1] Lewis, S. M., Asselin-Labat, M.-L., Nguyen, Q., Berthelet, J., Tan, X., Wimmer, V. C., Merino, D., Rogers, K. L., and Naik, S. H., “Spatial omics and multiplexed imaging to explore cancer biology,” Nat. Methods 18, 997–1012 (2021).

[2] Hickey, J. W., Neumann, E. K., Radtke, A. J., Camarillo, J. M., Beuschel, R. T., Albanese, A., McDonough, E., Hatler, J., Wiblin, A. E., Fisher, J., et al., “Spatial mapping of protein composition and tissue organization: a primer for multiplexed antibody-based imaging,” Nat. Methods 19, 284–295 (2022).

[3] Ali, H. R., Jackson, H. W., Zanotelli, V. R., Danenberg, E., Fischer, J. R., Bardwell, H., Provenzano, E., Rueda, O. M., Chin, S.-F., Aparicio, S., et al., “Imaging mass cytometry and multiplatform genomics define the phenogenomic landscape of breast cancer,” Nat. Cancer 1, 163–175 (2020).

[4] Jackson, H. W., Fischer, J. R., Zanotelli, V. R., et al., “The single-cell pathology landscape of breast cancer,” Nature 578, 615–620 (2020).

[5] Rendeiro, A. F., Ravichandran, H., Bram, Y., et al., “The spatial landscape of lung pathology during covid-19 progression,” Nature 593, 564–569 (2021).

[6] Lu, P., Oetjen, K. A., Bender, D. E., et al., “Imc-denoise: a content aware denoising pipeline to enhance imaging mass cytometry,” bioRxiv (2022).

[7] Goltsev, Y., Samusik, N., Kennedy-Darling, J., Bhate, S., Hale, M., Vazquez, G., Black, S., and Nolan, G. P., “Deep profiling of mouse splenic architecture with codex multiplexed imaging,” Cell 174, 968–981 (2018).

[8] Schürch, C. M., Bhate, S. S., Barlow, G. L., Phillips, D. J., Noti, L., Zlobec, I., Chu, P., Black, S., Demeter, J., McIlwain, D. R., et al., “Coordinated cellular neighborhoods orchestrate antitumoral immunity at the colorectal cancer invasive front,” Cell 182, 1341–1359 (2020).

[9] Lin, J.-R., Izar, B., Wang, S., Yapp, C., Mei, S., Shah, P. M., Santagata, S., and Sorger, P. K., “Highly multiplexed immunofluorescence imaging of human tissues and tumors using t-cycif and conventional optical microscopes,” Elife 7 (2018).

[10] Angelo, M., Bendall, S. C., Finck, R., Hale, M. B., Hitzman, C., Borowsky, A. D., Levenson, R. M., Lowe, J. B., Liu, S. D., Zhao, S., et al., “Multiplexed ion beam imaging of human breast tumors,” Nat. Medicine 20, 436–442 (2014).

[11] Keren, L., Bosse, M., Marquez, D., Angoshtari, R., Jain, S., Varma, S., Yang, S.-R., Kurian, A., Van Valen, D., West, R., et al., “A structured tumor-immune microenvironment in triple negative breast cancer revealed by multiplexed ion beam imaging,” Cell 174, 1373–1387 (2018).

[12] Zhou, J., Cui, G., Hu, S., Zhang, Z., Yang, C., Liu, Z., Wang, L., Li, C., and Sun, M., “Graph neural networks: A review of methods and applications,” AI Open 1, 57–81 (2020).

[13] Wu, Z., Pan, S., Chen, F., Long, G., Zhang, C., and Philip, S. Y., “A comprehensive survey on graph neural networks,” IEEE Trans. Neural Netw. Learn. Syst. 32, 4–24 (2020).

[14] Martin-Gonzalez, P., Crispin-Ortuzar, M., and Markowetz, F., “Predictive modelling of highly multiplexed tumour tissue images by graph neural networks,” in [Interpretability of Machine Intelligence in Medical Image Computing, and Topological Data Analysis and Its Applications for Medical Data], 98–107, Springer (2021).

[15] Innocenti, C., Zhang, Z., Selvaraj, B., Gaffney, I., Frangos, M., Cohen-Setton, J., Dillon, L. A., Surace, M. J., Pedrinaci, C., Hipp, J., et al., “An unsupervised graph embeddings approach to multiplex immunofluorescence image exploration,” bioRxiv (2021).

[16] Fischer, D. S., Schaar, A. C., and Theis, F. J., “Modeling intercellular communication in tissues using spatial graphs of cells,” Nat. Biotechnol., 1–5 (2022).

[17] Hu, J., Li, X., Coleman, K., et al., “Spagcn: Integrating gene expression, spatial location and histology to identify spatial domains and spatially variable genes by graph convolutional network,” Nat. methods 18, 1342–1351 (2021).

[18] Kim, J., Rustam, S., Mosquera, J. M., Randell, S. H., Shaykhiev, R., Rendeiro, A. F., and Elemento, O., “Unsupervised discovery of tissue architecture in multiplexed imaging,” Nat. Methods 19, 1653–1661 (2022).

[19] Brbić, M., Cao, K., Hickey, J. W., Tan, Y., Snyder, M. P., Nolan, G. P., and Leskovec, J., “Annotation of spatially resolved single-cell data with stellar,” Nat. Methods 19, 1411–1418 (2022).

[20] Wu, Z., Trevino, A. E., Wu, E., Swanson, K., Kim, H. J., D’Angio, H. B., Preska, R., Charville, G. W., Dalerba, P. D., Egloff, A. M., et al., “Graph deep learning for the characterization of tumour microenvironments from spatial protein profiles in tissue specimens,” Nat. Biomed. Eng 6, 1435–1448 (2022).

[21] Lu, M. Y., Williamson, D. F., Chen, T. Y., et al., “Data-efficient and weakly supervised computational pathology on whole-slide images,” Nat. Biomed. Eng. 5, 555–570 (2021).

[22] Hastie, T., Tibshirani, R., Friedman, J. H., and Friedman, J. H., [The elements of statistical learning: data mining, inference, and prediction], vol. 2, Springer (2009).

[23] Berg, S., Kutra, D., Kroeger, T., Straehle, C. N., Kausler, B. X., Haubold, C., Schiegg, M., Ales, J., Beier, T., Rudy, M., et al., “Ilastik: interactive machine learning for (bio) image analysis,” Nat. Methods 16, 1226–1232 (2019).

[24] Loi, S., Michiels, S., Salgado, R., Sirtaine, N., Jose, V., Fumagalli, D., Kellokumpu-Lehtinen, P.-L., Bono, P., Kataja, V., Desmedt, C., et al., “Tumor infiltrating lymphocytes are prognostic in triple negative breast cancer and predictive for trastuzumab benefit in early breast cancer: results from the finher trial,” Ann. Oncol. 25, 1544–1550 (2014).

[25] Levine, J. H., Simonds, E. F., Bendall, S. C., Davis, K. L., El-ad, D. A., Tadmor, M. D., Litvin, O., Fienberg, H. G., Jager, A., Zunder, E. R., et al., “Data-driven phenotypic dissection of aml reveals progenitor-like cells that correlate with prognosis,” Cell 162, 184–197 (2015).

[26] Traag, V. A., Waltman, L., and Van Eck, N. J., “From louvain to leiden: guaranteeing well-connected communities,” Sci. Rep. 9, 1–12 (2019).

[27] Kipf, T. N. and Welling, M., “Semi-supervised classification with graph convolutional networks,” arXiv preprint arXiv:1609.02907 (2016).

[28] Ba, J. L., Kiros, J. R., and Hinton, G. E., “Layer normalization,” arXiv preprint arXiv:1607.06450 (2016).

[29] Klambauer, G., Unterthiner, T., Mayr, A., and Hochreiter, S., “Self-normalizing neural networks,” Adv. Neural Inf. Process. Syst. 30 (2017).

[30] Berrada, L., Zisserman, A., and Kumar, M. P., “Smooth loss functions for deep top-k classification,” arXiv preprint arXiv:1802.07595 (2018).

[31] Paszke, A., Gross, S., Massa, F., Lerer, A., Bradbury, J., Chanan, G., Killeen, T., Lin, Z., Gimelshein, N., Antiga, L., et al., “Pytorch: An imperative style, high-performance deep learning library,” Adv. Neural Inf. Process. Syst. 32 (2019).

[32] Fey, M. and Lenssen, J. E., “Fast graph representation learning with pytorch geometric,” arXiv preprint arXiv:1903.02428 (2019).

[33] Pedregosa, F., Varoquaux, G., Gramfort, A., Michel, V., Thirion, B., Grisel, O., Blondel, M., Prettenhofer, P., Weiss, R., Dubourg, V., et al., “Scikit-learn: Machine learning in python,” J. Mach. Learn. Res. 12, 2825–2830(2011).

[34] Wolf, F. A., Angerer, P., and Theis, F. J., “Scanpy: large-scale single-cell gene expression data analysis,” Genome Biol. 19, 1–5 (2018).

[35] McInnes, L., Healy, J., and Melville, J., “Umap: Uniform manifold approximation and projection for dimension reduction,” arXiv preprint arXiv:1802.03426 (2018).

[36] Li, H., Calder, C. A., and Cressie, N., “Beyond moran’s i: testing for spatial dependence based on the spatial autoregressive model,” Geogr. Anal. 39, 357–375 (2007).

